# CoxBase: an online platform for epidemiological surveillance, visualization, analysis and typing of *Coxiella burnetii* genomic sequences

**DOI:** 10.1101/2020.11.29.402362

**Authors:** Akinyemi. M. Fasemore, Andrea Helbich, Mathias. C. Walter, Thomas Dandekar, Gilles Vergnaud, Konrad U. Förstner, Dimitrios Frangoulidis

## Abstract

Q (query) fever is an infectious zoonotic disease caused by the Gram-negative bacteria *Coxiella burnetii*. Although the disease has been studied since decades, it still represents a threat due to sporadic outbreaks across farms in Europe. The absence of a central platform for *Coxiella* typing data management in an important epidemiological gap which is relevant in the case of an outbreak. To fill this gap, we have designed and implemented an online, open-source, and, web-based platform called CoxBase (https://coxbase.q-gaps.de). This platform includes a database that holds genotyping information of more than 400 *Coxiella* isolates alongside metadata that annotates them. We have also implemented features for *in silico* genotyping of completely or minimally assembled *Coxiella* sequences using five different typing methods, querying existing isolates, visualization of isolate’s geodata via aggregation on a world map and submission of new isolates. We tested our *in silico* typing method on 50 *Coxiella* genomes downloaded from the RefSeq database and we successfully genotyped all except for cases where the sequence quality was poor. We identified new spacer sequences using our implementation of the MST *in silico* typing method, and established adaA gene phenotypes for all 50 genomes as well as their plasmid types.

## INTRODUCTION

Q (query) fever is an infectious zoonotic disease that affects humans and small ruminants like sheep, goat and cattle. It was first described among abattoir workers in Queensland, Australia, with symptoms of febrile illness in 1937 (1). The causative agent is a Gram-negative, pleomorphic, obligate intracellular bacterium called *Coxiella burnetii*. It has a world wide distribution and persists in biological and environmental reservoirs like milk, hay and dust, which can act as sources for sporadic outbreaks in livestock (2).

Since its first description as a febrile illness in Australia, the pathology of Q fever is now more understood and has been described as usually sub-clinical in ruminants but may manifest in form of late term abortion in pregnant ruminant females (3). In humans, the disease can be observed in two different forms. The first form is the acute disease, which is usually self-limiting, might occur alongside symptoms such as febrile illness, fever and severe headaches. It has been shown to happen in 40% of the primary Q fever cases. The second form is the chronic form, usually long lasting, characterized by endocarditis and can be severe and in dire cases fatal. It occurs in 1-5% of primary cases, the remaining cases are usually subclinical/asymptomatic and are also defined as acute disease (4) (2).

The epidemiology of this disease has been linked to the interplay of several dynamic factors including but not limited to, vector diversity, reservoir type and worldwide distribution of the disease (5). Another important point for disease control is the absence of a central platform that connects the different ends of the large and growing field of *Coxiella* research.

As a result, data from *Coxiella* research are dispersed over the academic space and if collected at a point is usually specific to a single method. The implication of these is that speed of research flow is significantly impeded especially in urgent cases of outbreaks where strain comparison and discrimination is vital to the control of the aetiological agent.

To highlight this challenge, there are up to five known genotyping methods for discriminating *Coxiella* species namely Multiple Locus Variable-number Tandem Repeat Analysis (MLVA) (6) (7), Multispacer typing (MST) (8), IS1111 typing (9), AdaA gene typing (10) and Plasmid typing (11) (12). MLVA and IS1111 typing require the measurement of PCR amplification products. MST requires the sequencing of intergenic regions whereas AdaA typing is based upon the sequencing of one coding sequence. All methods allow to detect a correlation between geographic origin and genotype and are useful for typing strains in endemic regions as well as clinical entities (5) (10). MLVA, MST and IS1111 methods offer a higher resolution compared to the other two methods (5).

A researcher interested in typing a new *Coxiella* strain is likely to employ more than a single method to obtain quality proof or at least to employ the methods accessible in his particular setting. Access to a database resource with strain information and metadata will be necessary for comparison purpose.

Presently there are two of such resources that house *Coxiella* genotyping data. The first is the MLVA databank (http://mlva.i2bc.paris-saclay.fr/mlvav4/genotyping/) and the second is the MST database (https://ifr48.timone.univ-mrs.fr/mst/coxiellaburnetii/), for the other genotyping methods there are no available database resources.

First we sought to overcome the lack of additional genotyping resource, then we sought to consolidate on the existing resources via introduction of new features such as visualization of allelic reference for MST typing, aggregation of MLVA groups and introduction of MLVA genotypes for better comparison. To this end, we have developed an online, open, web-based platform called CoxBase, which caters for vital aspects of internet based *Coxiella* research. This platform also includes a database that contains over 400 *C. burnetii* isolates from different countries. It has been implemented with a user interface for quick retrieval of isolate information as well as a submission channel to add to the growing body of new *Coxiella* isolates.

Also, we sought to unify all *Coxiella* typing systems under a single platform, alongside all the published details of *Coxiella* genotyping, including primers for genotyping protocols, as well as phenotypes, for the purpose of strain discovery and comparison. We implemented an *in silico* genotyping option for all major genotyping systems of *C*.*burnetii* based on whole genomic sequences.

Finally, we included visualization systems to quickly summarize all metadata on country level, maps for enhanced geographic localization of isolates and a worldwide distribution map of all *C. burnetii* isolates in our database. Here, we present our platform, its current scope, usage and capabilities.

## MATERIALS AND METHODS

### Webserver components

The server is run by Apache HTTP server (version 2.4.29), on a machine hosted by de.NBI Cloud services. The server components can be grouped under 2 main sections, the front end and the back end.

#### The front end

The main component of the front end is the web user interface, this is designed to accept user queries as well as submissions, send data to the back end and present data back to the user. Styling was achieved through an assortment of the cascading style sheet (CSS) Bootstrap framework (https://getbootstrap.com/), jQuery UI library and custom CSS style scripts. The validation of forms and events processing is achieved with JavaScript. The user interface accepts two kinds of data input: FASTA formatted whole genomic sequence (contigs or complete assembly) for typing purposes, and typing profiles via multi locus variable number tandem repeat analysis (MLVA) (7) and Multilocus sequence typing (MST) (8) for isolate comparison and discovery.

#### The back end

The back end handles user requests and uses a MySQL database to store data. Requests are handled via an Apache server (https://d.apache.org/) which then communicates via the Web Server Gateway Interface (WSGI) to a python pyramid framework application (https://trypyramid.com/). The application processes the request and communicates via the SQLAlchemy library (https://www.sqlalchemy.org) to the MySQL storage.

### Genome typing

We have implemented five different *in silico* typing methods for *Coxiella* sequences on the server. The MLVA typing method (6), the MST method (8), the adaA gene typing method (10), the plasmid typing method (12) and the 1S1111 typing method (9). The typing programs were implemented in the Python web application.

#### Establishing the typing features

#### MLVA typing

The MLVA typing feature accepts as input, genomic sequences either as contigs or complete assembly in FASTA format. The lengths of fourteen MLVA amplicons (when present) are extracted *in silico* with the e-PCR tool (13) using primers from Frangoulidis *et al*., 2014 (Table 1) (7) updated in http://mlva.i2bc.paris-saclay.fr/MLVAnet/spip.php?rubrique50. The repeat number is calculated with the formula below:

**Table 1.**
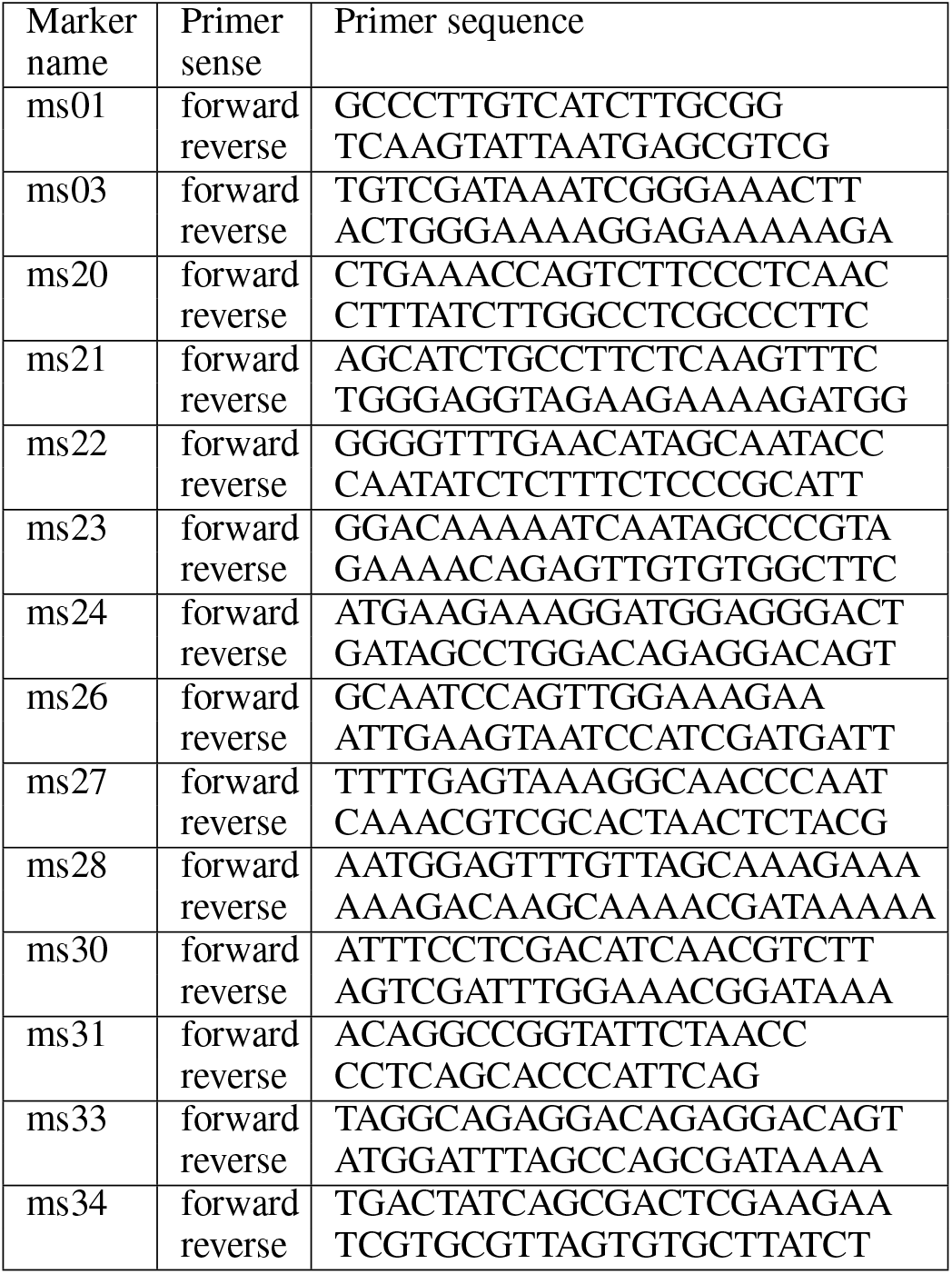
Table of used MLVA markers and their primer sequences

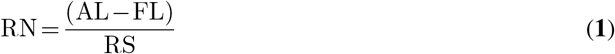

where RN = repeat number, AL = amplicon length, FL = flanking length and RS = repeat size.

For every submitted job, a unique identifier is generated that can be used to retrieve the results historically from the database within three weeks after the date of submission. The results of the MLVA typing are presented in form of a table with all the calculated parameters. A feature to search the database for closely related MLVA profiles is also provided.

#### MST typing

The *in silico* MST method accepts genomic sequence in FASTA format. The first step is amplicon detection via USEARCH (14). This is done using the MST primers from Glazunova *et al*., 2005 (8). The allele type is determined by aligning detected amplicon sequence globally with known alleles in the MST library (https://ifr48.timone.univ-mrs.fr/mst/coxiellaburnetii/spacers.html). Novel sequences with no match are also reported. The detected MST profile can be used as a query to the database to find the corresponding MST group.

#### IS1111 typing

The IS1111 typing is based on detection of localizations adjacent to IS1111 elements (9). This is a binary detection method meaning the discrimination is based on the absence or presence of an amplicon from a given location. For the *in silico* detection, we employed the e-PCR tool (13) to detect amplicons based on the primers described by (9) and extended by Bleichert & Hanczaruk 2012 (unpublished). The presence or absence is highlighted with a green (+) and red (-) respectively as shown in Figure 2.

**Figure 1.**
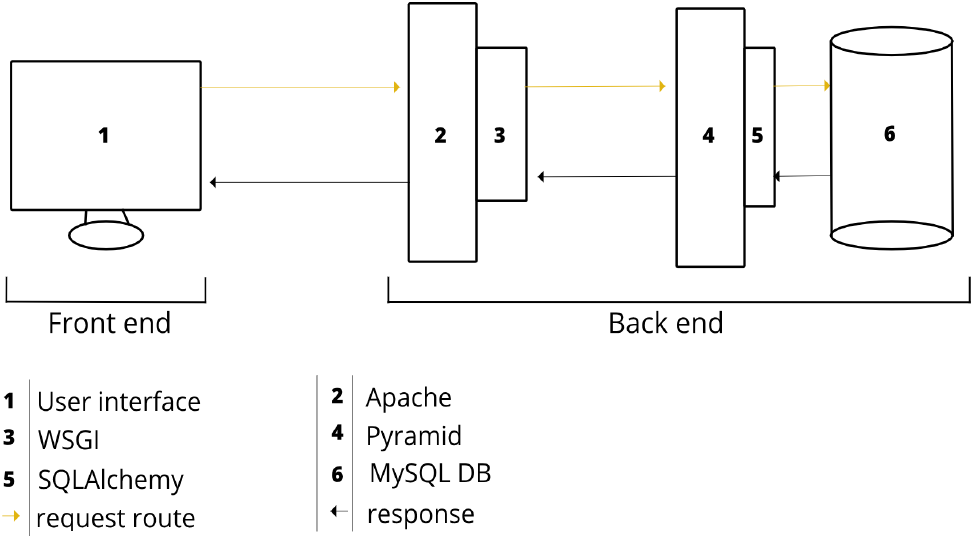
CoxBase Server Architecture.

**Figure 2.**
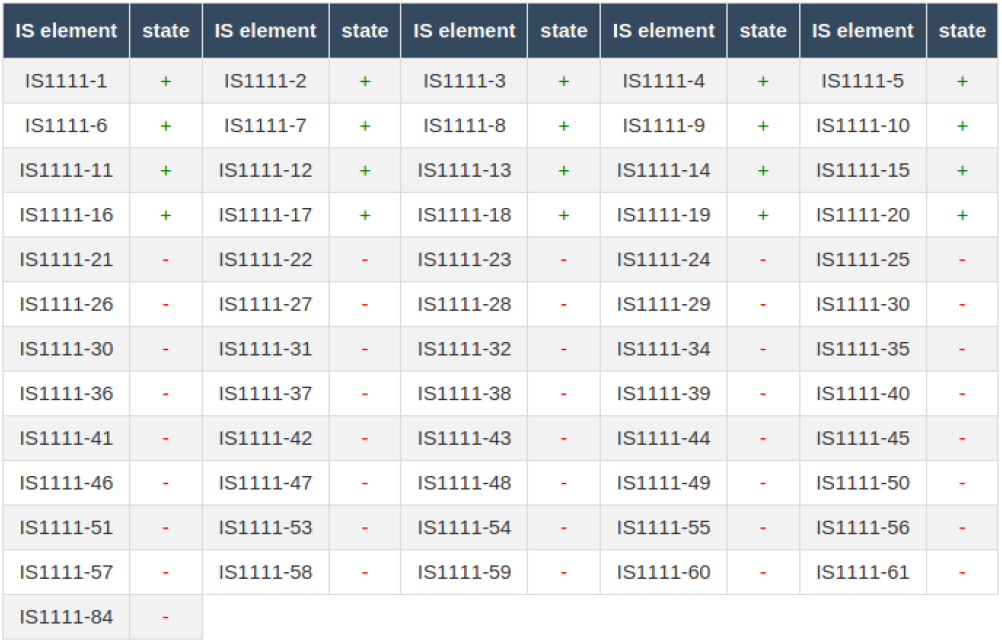
IS1111 typing result of RSA 439 as calculated on the CoxBase platform.

#### adaA and Plasmid typing

adaA phenotype has been earlier reported to correlate with plasmid type (10), therefore we combined these two typing methods together. Five different variants of the adaA gene have been reported, three single nucleotide variants (wildtype, A431T SNP, and repeat) and two deletion variants (Q154 deletion and Q212 deletion) (10). In our implementation, we first try to detect if the coding sequence of the adaA gene exists withing the genome to be typed. For this, we used the USEARCH tool (14) and the primer sequence for the detection of the entire adaA open reading frame (684 bases) (10). If an amplicon exits, we subsequently evaluate its length. If the length is longer than 684 bases then we assign it the adaA insertion genotype, and if it is shorter we assign it the incomplete adaA genotype. If it is exactly 684 bases, then we evaluate type of SNP at position 431 of the amplicon sequence. For the detection of plasmid type, we employed 4 primers that have been used for direct identification of *C. burnetii* plasmids via laboratory PCR methods (12) (11) (15).

### Isolate discovery and comparison

The CoxBase platform offers features for the discovery and comparison of *Coxiella* strains through several approaches. One approach is to query the database based on metadata and genotype features like country, host type, plasmid type, year of isolation, MLVA genotype and MST group. The advantage of this approach is that it’s fine grained and the fields can be aggregated to build more specific queries. Another approach utilizes a faceted search, this approach is more suitable for refining queries based on reviewed criteria. Other approaches rely on making queries based on known typing profiles via MLVA or MST typing schema. This is implemented as follows:

For a user who wishes to discover isolates with a specific isolate profile (MST or MLVA). They need to provide a complete or partial profile (MLVA or MST) of the isolate which they are interested in. Usually one marker is enough for a search but for a more defined and reliable results, at least 6 markers should be provided for the MLVA query and 10 for the MST. For ease of comparison, isolates with similar profiles are pooled together in a single row in the query results. Profile entries can then be expanded with the click of a button called View profile entries in the final column of the result table. A list of all isolates with that profile is provided with metadata. Geographical information is visualized through a Leaflet Map (https://leafletjs.com/). The aim is to provide a geographic orientation that can be used to estimate physical proximity of the isolates. Figure 3 shows a single isolate from the A14 MLVA group that was isolated from a sheep in Hannover, Germany. The last approach relies on grouping based on geographical location. A user interested in isolates from a particular country will approach the distribution map. He can access a comprehensive table of all isolates from the country of interest after clicking on the country marker located on the map.

**Figure 3.**
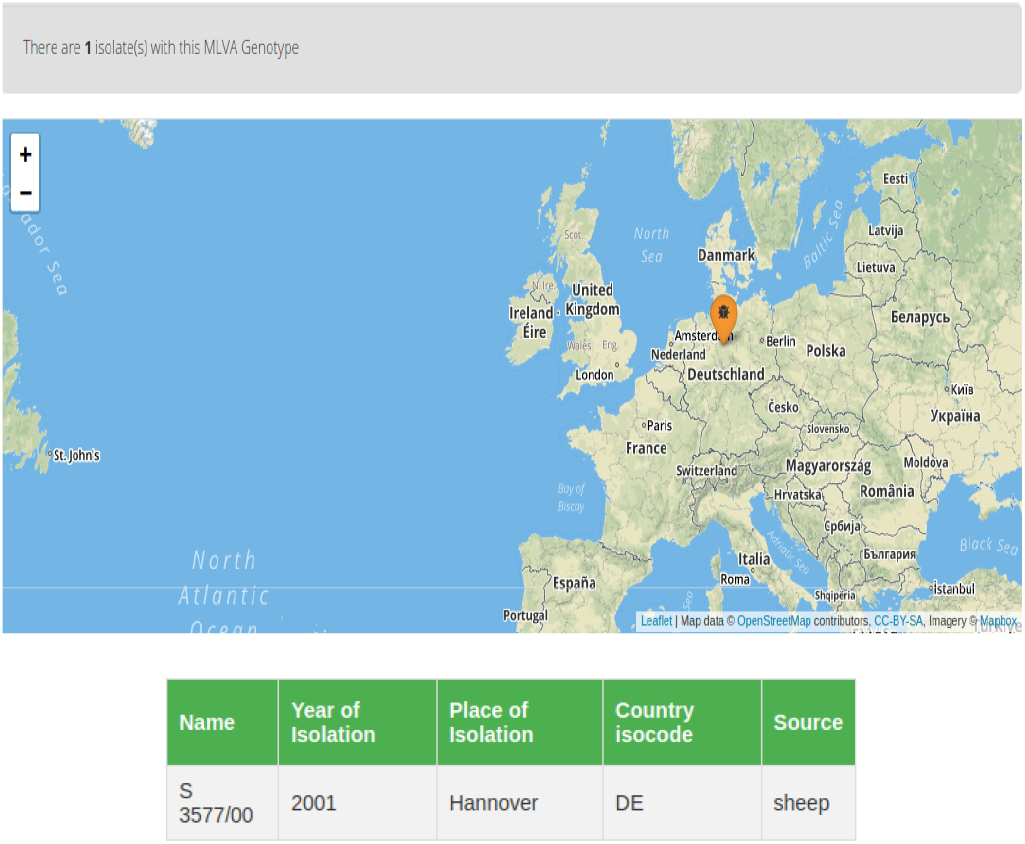
A strain from the A14 MLVA group that was isolated from a sheep in Hannover, Germany.

### Visualization

We implemented an interactive visualization feature based on the Chart.js (https://www.chartjs.org/) JavaScript visualization library. This can be accessed through the dashboard link on country markers in the distribution map. Distribution plots for metadata categories such as host type, year of isolation, place of isolation as well as genotype could help answer questions such as the most predominant host type in a particular location as illustrated in Figure 4.

**Figure 4.**
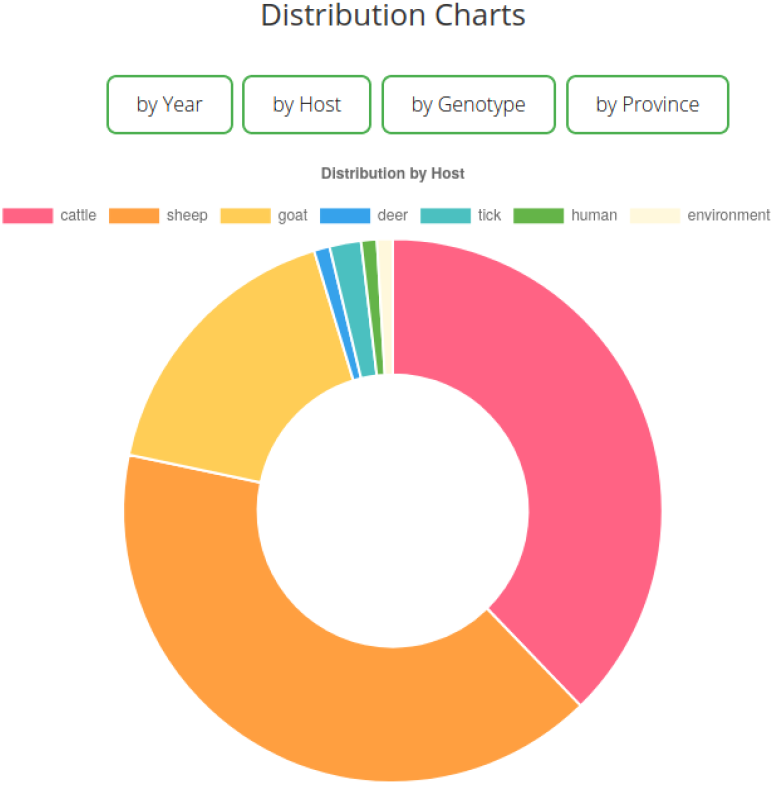
Donut plot of host data from Germany and it shows that most common hosts are sheep and cattle.

**Figure 5.**
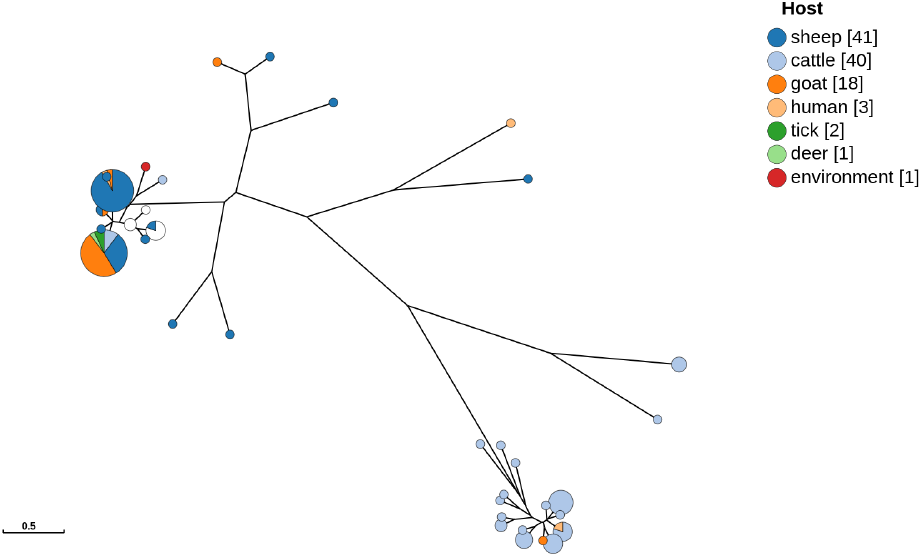
GrapeTree visualization of *C. burnetii* isolates from Germany on CoxBase based on MLVA genotyping. Distinctive clusters based on metadata such as host type can be inferred from such a tree.

**Figure 6.**
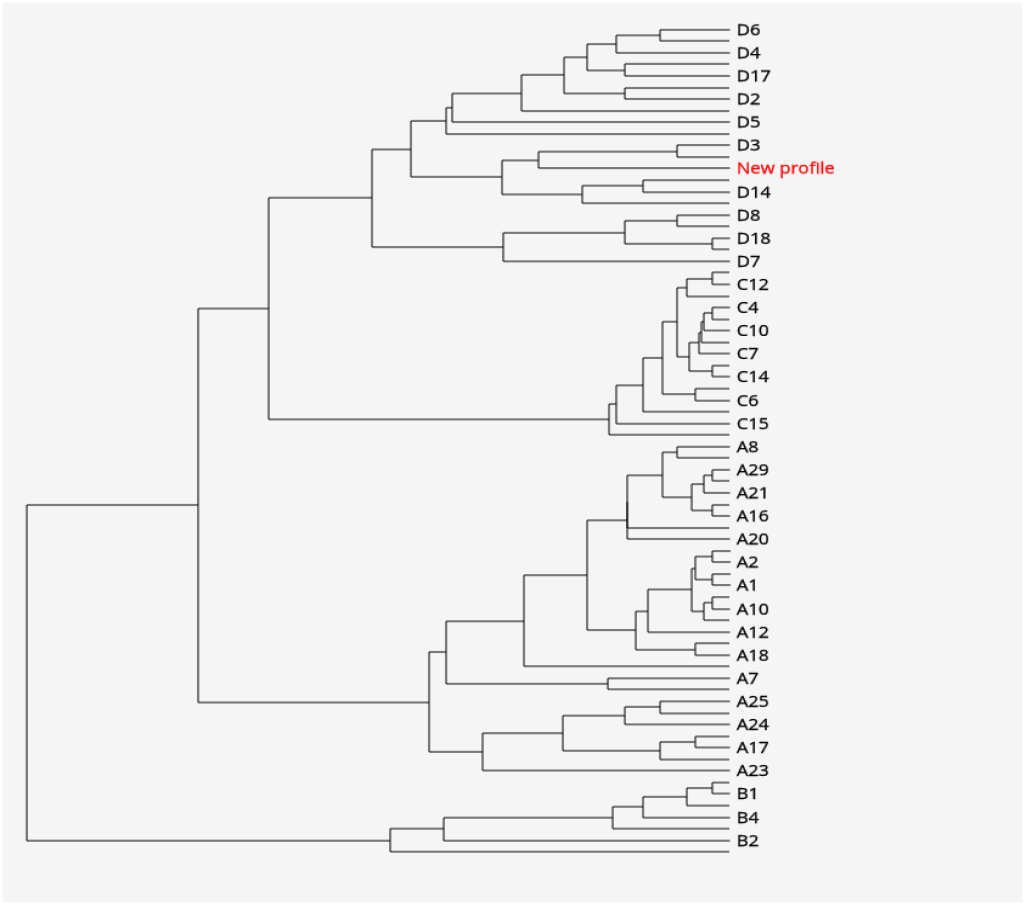
Unrooted phylogenetic tree of all MLVA genotypes. Highlighted node shows the position of *C. burnetii* strain Q321 that was isolated from a cow’s milk in Russia. MLVA typing was done via CoxBase.

### Data collection sets and Testing

We tested our implementation on fifty *Coxiella* whole genomic sequences from RefSeq. (Table of genomes in supplementary table). We had eleven complete chromosome assembly, thirteen chromosome assembly, fifteen contigs and eleven scaffolds. The average genome size was 2.01Mb. Forty of these were of unknown genotypes (not published if known), so we sought to type these genomes as well. The genome sequences in FASTA format were downloaded from the RefSeq database and stored without any form of modification. The genomes were typed individually using the different typing methods on our platform. After which the results were compared to known strains in our database. The results are discussed below

## RESULTS: APPLICATION EXAMPLES

### MLVA typing

Based on the *in silico* typing scheme we could separate the fifty genomes under thirteen MLVA groups. We observed that some markers were often not detected *in silico*. We computed the effectiveness for each marker (The probability of the marker producing an amplicon *in silico* as a percentage) based on the 50 *Coxiella* genomes. We observed that six of the fourteen markers had a marker effectiveness of one hundred percent, six markers between ninety-eight to seventy-two percent effectiveness and two markers with an effectiveness of fifty-six and twenty percent (Table of Marker effectiveness in Supplementary material). For genomes with known MLVA types (RSA 493 and Dugway), we were able to classify them into the correct MLVA group as described already in Frangoulidis *et al*., (2014) (7).

### MST typing

We were able to classify the fifty genomes into twelve MST groups, except for two genomes whose profile had no matching MST group and one with an undetermined spacer, therefore it could not be placed in an appropriate MST group. We were able to confirm the MST group of genomes with published MST genotypes (Ohio, Henzerling, and Nine mile), however, our results differ from the published MST genotype of the Dugway strain. The spacer profile indicated it belonged to the MST type 60 and not MST 20 as published. Nearly half of the genomes (n=24/50) are either MST 16 or MST 61. All the MST primers had an amplicon *in silico*. We also detected six new spacer alleles in seven genomes. The MST markers were reliable. However, we observed that detection of a new spacer allele might be a false positive due to incomplete sequences in the alignment. Therefore, we implemented an alignment visualization feature for the detection of new spacer sequences based on the BlasterJS library (16).

### AdaA and Plasmid typing

The plasmid of the twenty-three genomes out of the fifty genomes in the dataset were already known. We were able to confirm the known plasmid type and extended the information with their adaA gene phenotypes, and for genomes whose plasmid type was not known we were able to detect the plasmid type as well as their adaA gene phenotypes. We determined 43 out of the 50 genomes carried a plasmid. QpH1 was present in 34 instances. Nine out of the fifty genomes had an adaA deletion phenotype and the remaining were adaA wildtype. Out of the 41 positive strains, seven had a repeat insertion in there adaA gene sequence, while four genomes showed a SNP at position 431 of the adaA gene sequences. We observed that six out of the seven genomes with an adaA repetition phenotype belonged to the same MST profile.

### Phylogenetic analysis

We implemented two types of visualization for phylogenetic trees. The first tree is a GrapeTree (17) implementation that can be used to visualize genomic relationships of grouped data based on their MLVA profiles. The resulting tree can be color coded based on metadata, editable and can also be exported into several image formats. The second tree is implemented using the PhyD3 visualization library (18). This is especially useful for locating MLVA profile in the MLVA genotype tree, thereby associating a strain with a new MLVA profile with its closest MLVA genotype.

## DISCUSSION

Here we present a platform that was built with the aim to overcome the lack of a centralized genomic data resource for *Coxiella burnetii*.

This is the first genotyping platform that combines all the disparate typing systems of *Coxiella burnetii*. Similar platforms exist for other bacteria species such as PubMLST albeit usually focused on a single typing system.

Several features are particularly novel and unique: We combined five typing methods to enable rapid identification of *Coxiella* strains as well as the visualization of the metadata coupled to the geographical distribution. The latter format is particularly useful to study and control outbreaks, the major shortcoming for which our platform was constructed.

We have also included several features that could assist researchers to understand the variability within the genomes of *C. burnetii* in an epidemiological context. We have leveraged on technologies such as NGS, cloud computing and database to create an open web resource that can be used to genotype draft or completely assembled *C. burnetii* genomic sequences as well as compare them to existing strains. Our approach also brought together different aspects of *Coxiella* research including epidemiological surveillance, sequence analyses and phylogeny under a single platform. The strength of *in silico* typing methods rely on, to a significant degree, the quality of the input sequence. We observed that certain MLVA markers are also not as effective as others, however, a combination of multiple markers of different typing schemes should improve the averaging effect. Nevertheless, our implementations suggest that *in silico* typing can be an indispensable tool for rapid genotyping of *Coxiella* genomic sequences. We tested the implementation on 50 *C. burnetii* genomes from NCBI and we were able to type all except for cases where the sequence quality was not good enough. We could observe perfect corroboration with known genotypes when we used our implementation to type these sequences except for one case where we argue that the published profile might not be correct as the observed spacer profile differed in all alleles compared to the published profile. One limitation of our method is in the adaA gene typing. Although we can distinguish between the different adaA gene positive variants, we are yet to implement a feature to differentiate between the deletion variants (if it is a Q212 deletion or Q514 deletion). For now we only report if the adaA gene deletion exists in a given sequence and not the variant of the deletion type. We implemented a retrieval feature on CoxBase that will enable researchers to access the results of their typing analyses up to three weeks after their submission date. This would ease collaboration efforts on typing projects and reduce the complexity of information sharing. We have also implemented a genome browser for sequence visualization to accompany sequence typing investigations most especially primer analysis. Finally, we implemented a submission feature for researchers who wish to share new MLVA or MST profiles. We hope this platform will provide researchers with the opportunity to investigate the variability between *C. burnetii* genomes as well as help to better understand the epidemiology of the Q fever disease in terms of genotype correlations with metadata like host specificity and geographical information. We will update the platform periodically to keep the data current and curated.

## ACKNOWLEDGEMENTS

This research has been funded by the Federal Ministry of Education and Research of Germany (BMBF, project number 01KI1726E). Furthermore, the work was supported by the BMBF-funded de.NBI Cloud within the German Network for Bioinformatics Infrastructure (de.NBI) (031A537B, 031A533A, 031A538A, 031A533B, 031A535A, 031A537C, 031A534A, 031A532B).

## Conflict of interest statement

None declared.

## Notes

### Competing Interest Statement

The authors have declared no competing interest.

